# Promoter proximal pausing limits Yki-induced tumorous growth in *Drosophila*

**DOI:** 10.1101/813022

**Authors:** Sanket Nagarkar, Ruchi Wasnik, Pravallika Govada, Stephen Cohen, LS Shashidhara

## Abstract

Promoter proximal pausing (PPP) of RNA Polymerase II has emerged as a crucial rate-limiting-step in the regulation of gene expression. Regulation of PPP is brought about by complexes 7SK snRNP, P-TEFb (Cdk9/cycT) and the Negative Elongation Factor (NELF) which are highly conserved from *Drosophila* to humans. Here we show that RNAi-mediated depletion of *bin3* or *Hexim* of the 7SK snRNP complex or depletion of individual components of the NELF complex enhance Yki-driven growth leading to neoplastic transformation of *Drosophila* wing imaginal discs. We also show that increased CDK9 expression cooperates with Yki, in driving neoplastic growth. Interestingly, over-expression of CDK9 on its own or in the background of depletion of one of the components of 7SK snRNP or the NELF complex necessarily and specifically needed Yki over-expression to cause tumorous growth. Genome-wide gene expression analyses suggested that deregulation of protein homeostasis is associated with tumorous growth of wing imaginal discs. As both Fat/Hippo/Yki pathway and PPP are highly conserved, our observations may provide insights into mechanisms of oncogenic function of YAP, the orthologue of Yki in human.

## Introduction

Regulation of growth is arguably the most critical phenomenon that establishes size and shape of all tissues, organs and overall body size in metazoan animals. It is also an important homeostatic process, failure of which is linked to diseases and disorders, particularly cancer in human. Regulated growth is achieved by an intricate interplay between factors promoting growth (oncogenes) and those suppressing it (tumor suppressors).

Yorkie (Yki), the *Drosophila* ortholog of the Yes-Associated Protein 1 (YAP1), acts as a transcriptional cofactor that mediate the effects of the Hippo tumor suppressor pathway. The Hippo pathway is highly conserved from *Drosophila* to humans[1]. The Hippo (Hpo)/MST kinases and the Warts (Wts)/LATS kinases and their cofactors form kinase cassettes that directly phosphorylate Yki (YAP/TAZ) to regulate protein stability and activity[2]. Members of this pathway were initially found to limit tissue growth in *Drosophila* by limiting Yki activity [3,4]. Consistent with this YAP overexpression has been reported as a driver of tissue growth and cancer in a mouse model [4,5]. In humans, the YAP1 locus was found to be amplified in different types of cancer [6,7]. These findings have sparked a great deal of interest in understanding of regulation of Yki/YAP function.

In *Drosophila*, Yki regulates expression of regulators of cell growth and survival such as *Diap1, dMyc, bantam* etc. Targets of YAP in humans include the EGFR-ligand AREG as well as *CTGF, Cyr61* [8]. While these target genes are necessary for growth induced by Yki/YAP activity, they are not sufficient to phenocopy effects of Yki/YAP. This indicates possibility of more regulators that are involved in Yki/YAP induced growth.

We have reported an *in vivo* screen in *Drosophila* [9] wherein we have identified a large number of genes which, when depleted, enhanced growth induced by Yki and EGFR. More importantly, these genes function like classical tumor suppressors as when down regulated in the background of over-expressed Yki or EGFR, we observed neoplastic growth. Among these, we identified a number of genes involved in the control of promoter proximal transcriptional pausing.

Promoter proximal pausing (PPP) of RNA Pol II has been identified as a key step in transcriptional regulation for many genes, during development and in stem cells [10–12]. At paused loci, after initiation, the RNA Pol II first passes through the promoter but then stops at about 30-60bp from transcription start site [13]. Productive transcription then requires release from the paused state. PPP is brought about by the Negative transcription elongation factor (NELF) and DRB-sensitivity inducing factor (DSIF) protein complexes, which were identified as factors responsible for DRB sensitivity of transcription elongation [14,15]. These complexes bind RNA Pol II and halt its progress downstream of the promoter. This pause is alleviated by a positive transcription elongation factor complex (P-TEFb) (Fig.1A), which consists of Cyclin T and Cyclin Dependent Kinase- CDK9 [16]. Once recruited to the paused complex, CDK9 phosphorylates NELF and DSIF leading to ejection of NELF from the paused complex while DSIF assists Ser-5 phosphorylated RNA Pol II in productive elongation [17]. The PTEFb complex is in turn regulated through sequestration by 7SK snRNP complex. P-TEFb is required for release paused RNA polymerase II into productive elongation [13]. Sequestration of P-TEFb by 7SK snRNP leads to its unavailability for mediating pause release, which in turn regulates transcription elongation via sustained pause of RNA Pol II. Mammalian 7sk-snRNP complex consists of 7sk RNA, Hexim1/2, Larp7 and MePCE. *Drosophila* homologs of components of mammalian 7sk-snRNP complex were identified and characterized recently [18]. These include -Bin3 (MePCE ortholog), Larp (Larp7 ortholog), Hexim (HEXIM1/2 ortholog) and d7SK RNA. These were shown to be conserved with the mammalian counterparts functionally. Liberation of P-TEFb is triggered by signaling events of pathways such as ERK, TCR etc. Thus, making sequestration and liberation of P-TEFb a context dependent process which is critical for regulating expression of gene regulation depending on the context.

**Figure 1.**
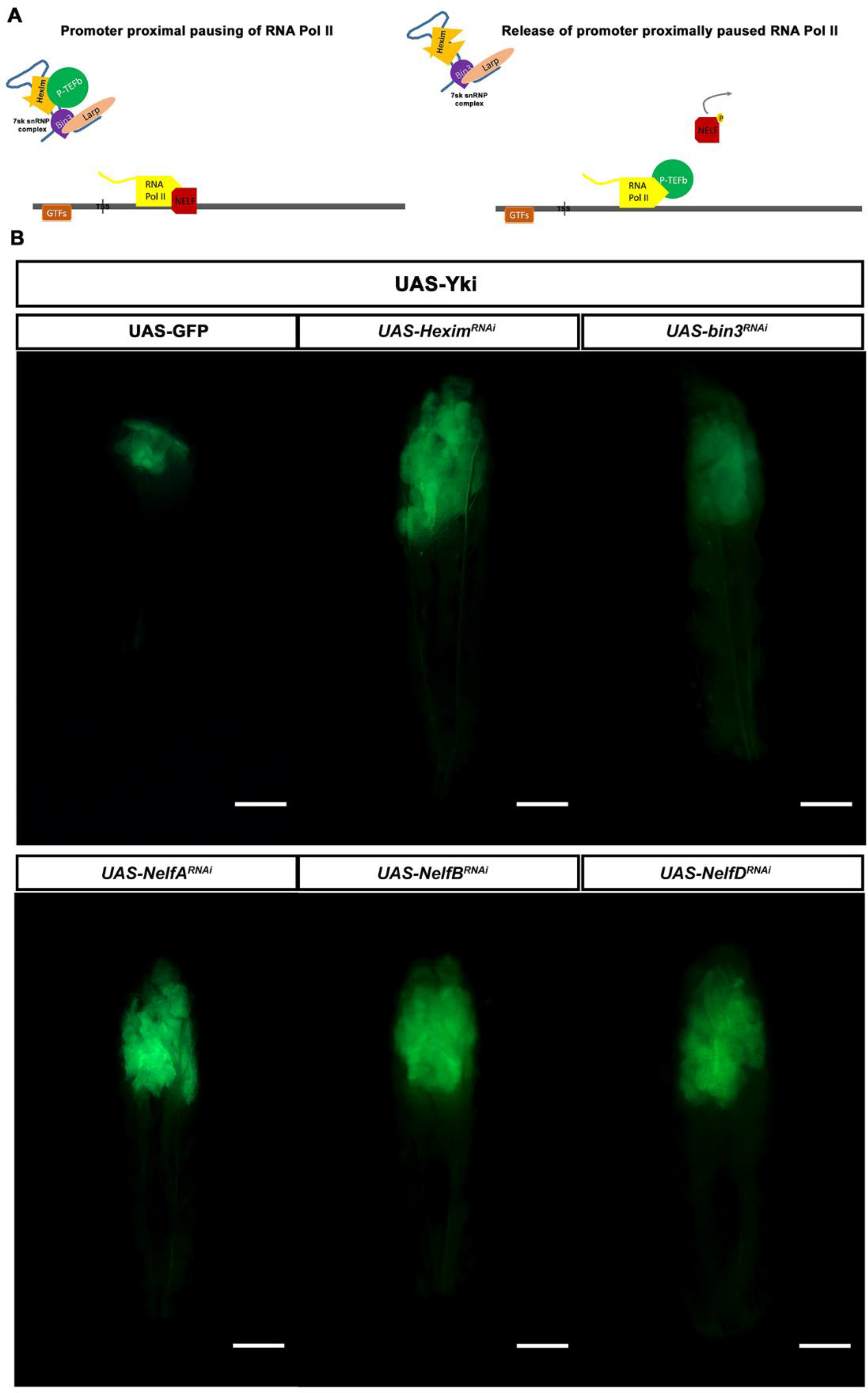
Identification of complexes invovled in promoter proximal pausing as tumor suppressors. A. schematic representing known function of two complexes we identified as candidate tumor suppressors. 7SK snRNP complex regulates promoter proximal pausing by sequestring P-TEFb complex, while NELF complex is invovled in the formation of stall of RNA Pol II at promoter proximal region. As dectated by surrounding cues, P-TEFb is released by 7SK snRNP comlex. Thus, freed P-TEFb is recruited to stalled RNA Pol II where it brings about release of RNA Pol II from the paused state. B. Larval images showing wing imaginal discs expressing GFP at low magnification. Dimensions of GFP-expressing tissue is indication of growth im imaginal discs. Top row - Larvae over-expressing only Yki (crossed to UAS-GFP as control) and those in combination with RNAi-mediated knock down of 7sksnRNP components- *Hexim* or *bin3* using GAL80^TS^; *ap*-GAL4; UAS-GFP (scale bar = 1 inch). Bottom row - wing discs over-expressing Yki in combination with RNAi-mediated knock down of of NELF components (from left to right)- *NelfA*, *NelfB*, *NelfD* (also known as TH1) using GAL80^TS^; *ap*-GAL4; UAS-GFP. Note significantly larger GFP expressing-wing tissue (green) in larvae that are over-expressing Yki and also depleted for a component of PPP.

Interestingly, CDK9 has been shown to be important for transcription of target genes of oncogenes such as Myc [19] and YAP [20]. Here, we present evidence of tumor suppressor function of various components involved in PPP, specifically in the context of elevated Yki activity. Our findings show that factors involved in PPP and its regulation are important to restrict Yki driven growth and to prevent neoplastic transformation *in vivo*.

## Results

### Depletion 7SK snRNP complex components cooperates with Yki in causing tumorous growth

Studies using *Drosophila* tumor models have found that larvae containing proliferating tumors are unable to enter pupariation and continue to grow [21]. The resulting giant larva phenotype can be used in genetic screens to identify tumor causing genotypes. We made use of this property to identify candidate genes in a genetic screen for tumor suppressors cooperating with Yki (the entire screen is published elsewhere [9]). We found that RNAi-mediated depletion of *bin3* and *Hexim*, components of the 7SK snRNA complex in combination with Yki overexpression led to massive overgrowth in wing disc tissue and giant larval phenotype (Fig.1B). Wing discs expressing Yki alone show only moderate overgrowth phenotype, and larvae eventually pupate (Fig 1B). Depletion of 7SK snRNP components did not produce overgrowth on their own (Fig.EV1), but only did so when coupled with Yki overexpression. We also did not observe wing disc overgrowth when depletion of 7SK snRNP components in combination with overexpression of other well-known onco-proteins such as EGFR or Notch intracellular domain (NICD) (data not shown). Thus, our observations suggest that, *Drosophila* 7SK snRNP complex may function to repress tumorigenic potential of Yki *in vivo* in an epithelial tumor model.

### Components of the NELF complex may function as tumor suppressors

The NELF complex is composed of four sub-units- NELF-A, B, C/D and E. Depletion of each NELF component using RNAi in combination with Yki also produced a giant larval phenotype (Fig. 1B) and massively overgrown wing disc tissue compared to the larvae only over-expressing Yki (Fig. 1B). Depletion of the NELF components on their own did not cause such giant larval phenotype or overgrowth of the wing disc tissue (Fig. EV1). These components too did not show any tumor phenotype in the context of over-expressed EGFR or NICD (data not shown).

It was intriguing to find multiple components of the two spatio-temporally separated protein complexes, involved in regulation of transcription elongation, among the tumor suppressors identified in a genome wide screen for factors cooperating with Yki in growth regulation [9].

### Neoplastic transformation induced by Yki combined with depletion of 7SK snRNP or NELF complexes

Yki is known to promote cell proliferation and cell survival. Thus, it is possible that larger size of the wing disc tissue observed upon loss of either 7SK snRNP or NELF complex is a result of enhancement of growth and survival effect of Yki and not a neoplastic transformation. To distinguish between the two possibilities, we analyzed the tumor tissue using markers that indicate neoplastic transformation.

First, we examined epithelial cell polarity. Neoplastic transformation of an epithelial tissue is accompanied by the loss of their characteristic apico-basal cell polarity. E-cadherin (E-Cad) is a sub-apically localized protein which provides a convenient marker for epithelial polarization [22]. Wing discs overexpressing Yki alone showed localization of E-Cad, in a pattern similar to the wild type wing discs, although former discs are much larger (Fig. 2A). This indicated that Yki overexpression caused overgrowth of the epithelium without perturbation of epithelial cell polarity. In contrast, when Yki overexpression was combined with depletion of a component of the 7SK snRNP complex or the NELF complex, sub-apical localization of E-cad was lost or perturbed (Fig.2A). Additionally, we analyzed F-actin, which localizes near the apical junctions of the wing disc epithelial cells, using Rhodamine labelled Phalloidin. As with E-Cad, we observed loss of apical localization of F-Actin in the Yki expressing tissue depleted of a component of the 7SK snRNP or the NELF complex, but not in wing disc tissue expressing Yki alone (Fig EV2). We did not observe any change in cell polarity as indicated by E-Cad and F-actin localization in wing discs with depletion of components of 7SK snRNP and NELF complexes alone (data not shown).

**Figure 2.**
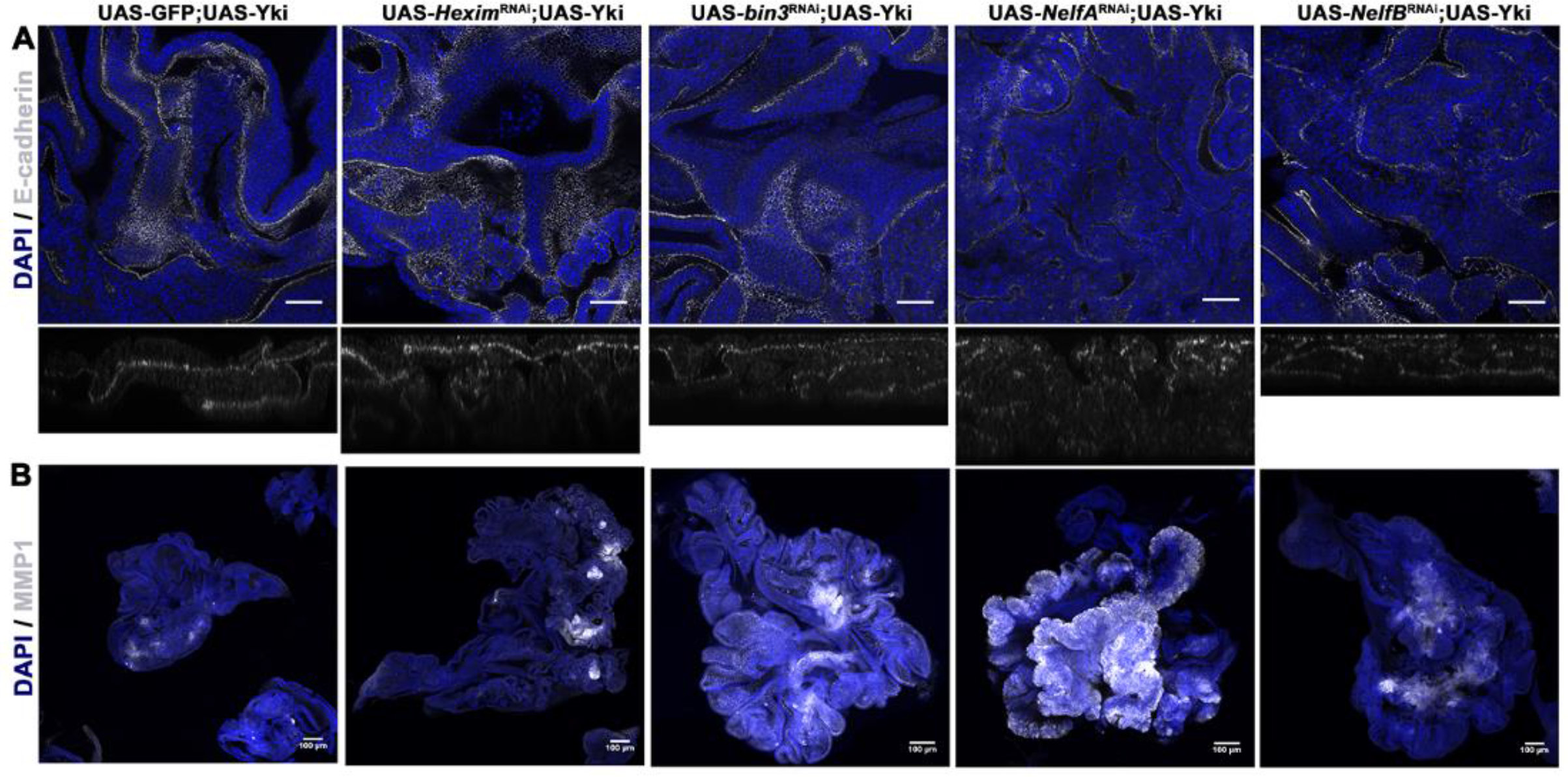
Characterization of tumors induced in the wing disc. A. Disruption of characteristic epithelial apico-basal polarity in tumor discs. Images of wing discs over-expressing Yki alone (crossed to UAS-GFP as control) or in combination RNAi-mediated knowckdown of *Hexim, bin3, NelfA* or *NelfB* using GAL80^TS^; *ap*-GAL4; UAS-GFP (scale bar = 10μm). Discs are stained for E-cad (white) expression and localization. Bottom panel of each image shows orthogonal optical section of respective genotype. Note delocalization of E-cad in tumorous tissues caused by the depletion of a component of PPP and Yki over-expression. All discs are also stained with DAPI (blue) to visualize nuclei. B. Increased expression of MMP1 is observed in tumor discs. Images of wing discs over-expressing Yki alone (crossed to UAS-GFP as control) or in combination RNAi-mediated knowckdown of *Hexim, bin3, NelfA* or *NelfB* using GAL80^TS^; *ap*-GAL4; UAS-GFP (scale bar = 100μm). Wing disc tissues are stained for MMP1 (white). Note increased MMP1 staining in tumorous tissues caused by the depletion of a component of PPP and Yki over-expression. All discs are also stained with DAPI (blue) to visualize nuclei.

The matrix metallo-protease MMP1 has been used as a marker of epithelial to mesenchymal transition (EMT) and neoplastic transformation in *Drosophila* tumor models. MMP1expression is elevated in tumor models and its depletion by RNAi has been reported to block metastasis [23–25]. We examined the effects of depleting components of 7SK snRNP and NELF complexes in Yki-expressing tissue on the levels of MMP1 by immunohistochemistry. We observed elevated levels of MMP1 in wing discs over-expressing Yki and depleted for Bin3, Hexim or the NELF complex (Fig 2B). We did not observe such increase in MMP1 levels in wing discs expressing Yki alone (Fig. 2B), neither in wing discs in with depletion of components of 7SK snRNP and NELF complexes alone (data not shown).

Taken together, tumors formed upon depletion of 7SK snRNP or NELF complex components in combination with Yki exhibit neoplastic characters. As neither genetic change alone produced these results, it appears that they act in combination to promote neoplasia, a classical mechanism of cooperative tumorigenesis as known in mammals. These observations provide evidence that the activity of 7SK snRNP and NELF complexes may have a tumor suppressing function, but only in the context of elevated Yki activity.

### CDK9 is required for Yki mediated tumorigenesis

The 7SK snRNP and NELF complexes, help in maintaining the paused state of RNA Pol II. Our findings raised the question of whether pausing of RNA Pol II per se served to limit the tumor promoting potential of Yki activity. If this is the case, we reasoned that using an alternative means to release RNA Pol II should also lead to tumorigenesis in the context of Yki overexpression. The P-TEFb complex, comprising of cycT/CDK9, is required for release of paused RNA Pol II and effective elongation of mRNA. CDK9 phosphorylates NELF complex leading to eviction of NELF from the pause site. This in turn facilitates release of paused RNA Pol II aiding in productive elongation. CDK9 also acts directly on RNA Pol II, phosphorylating it on S5 in the C-terminal domain, a known mark of elongating RNA Pol II [26]. As the P-TEFb complex is normally rendered inactive through sequestration by 7SK snRNP complex, we hypothesized that overexpressing CDK9 might bypass normal regulation of pausing, leading to inactivation of NELF complex and RNA Pol II release. Consistent with this hypothesis, we indeed observed massive tissue overgrowth when Yki was co-expressed with CDK9, while over-expression of CDK9 alone did not cause any such phenotype (Fig 3A). Wing discs expressing UAS-CDK9 together with UAS-Yki also showed loss of apically localized E-Cad as well as elevated MMP1 expression, compared to tissue expressing UAS-CDK9 alone or UAS-Yki alone. This indicates neoplastic transformation in wing discs co-expressing Yki and CDK9, similar to the transformation caused by depletion of 7SK snRNP and NELF complex components in combination with over-expressed Yki (Fig 3B).

**Figure 3.**
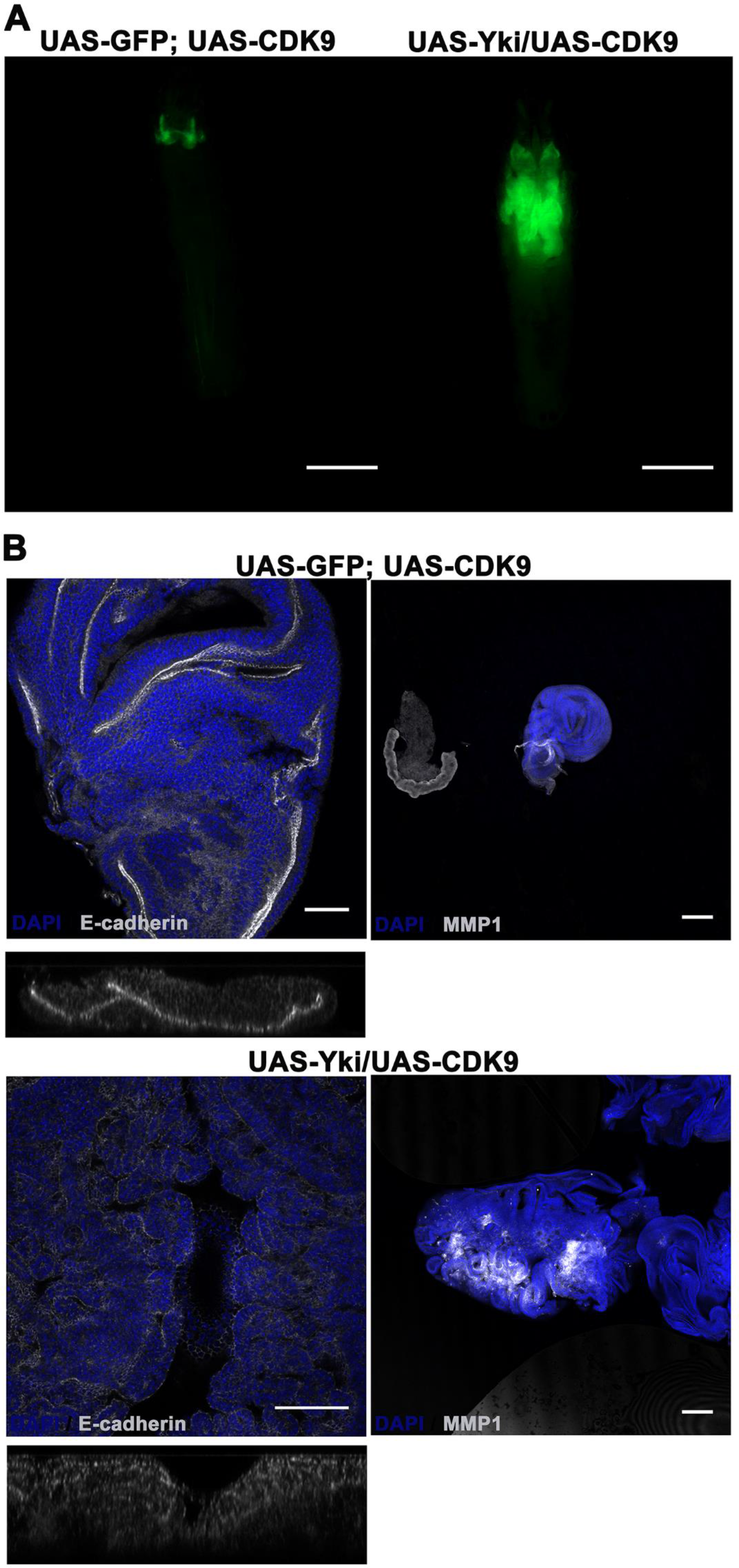
CDK9 cooperates with Yki in tumorigenesis. A. Larval images showing growth observed in combination of UAS-CDK9 and UAS-Yki as compared to UAS-CDK9 alone (crossed to UAS-GFP as control) using GAL80^TS^; *ap*-GAL4; UAS-GFP (scale bar = 1 inch). The combined over-expression phenocopies the phenotypes observed in Fig. 2B. B. Characterization of tumor tissue caused by combined over-expression of CDK9 and Yki using GAL80^TS^; *ap*-GAL4. Top row of images shows wing disc tissue over-expressing CDK9 alone, while the bottom row shows combined over-expression of CDK9 and Yki. Discs in the left column are stained for E-cadherin (white) (scale bar = 10μm) and in the right column are stained for MMP1 (white) expression (scale bar = 100μm). Please note deregulated E-cad localization (see in optical z-section given below the discs) and increased MMP1 expression in tissues which over-express both CDK9 and Yki, suggesting their neoplastic tumor state. All discs are also stained with DAPI (blue) to visualize nuclei.

As further test of this model, we asked whether CDK9 is essential for tumorigenic cooperation between depletion of 7SK snRNP complex components and Yki. Depletion of *cdk9* effectively suppressed the tissue overgrowth caused by depleting *bin3* or *Hexim* in Yki expressing tissue (Fig 4A). Those wing discs also showed normal apical localization of E-Cad and wildtype levels of MMP1 expression, suggesting complete suppression of tumorous growth (Fig. 4B, C).

**Figure 4.**
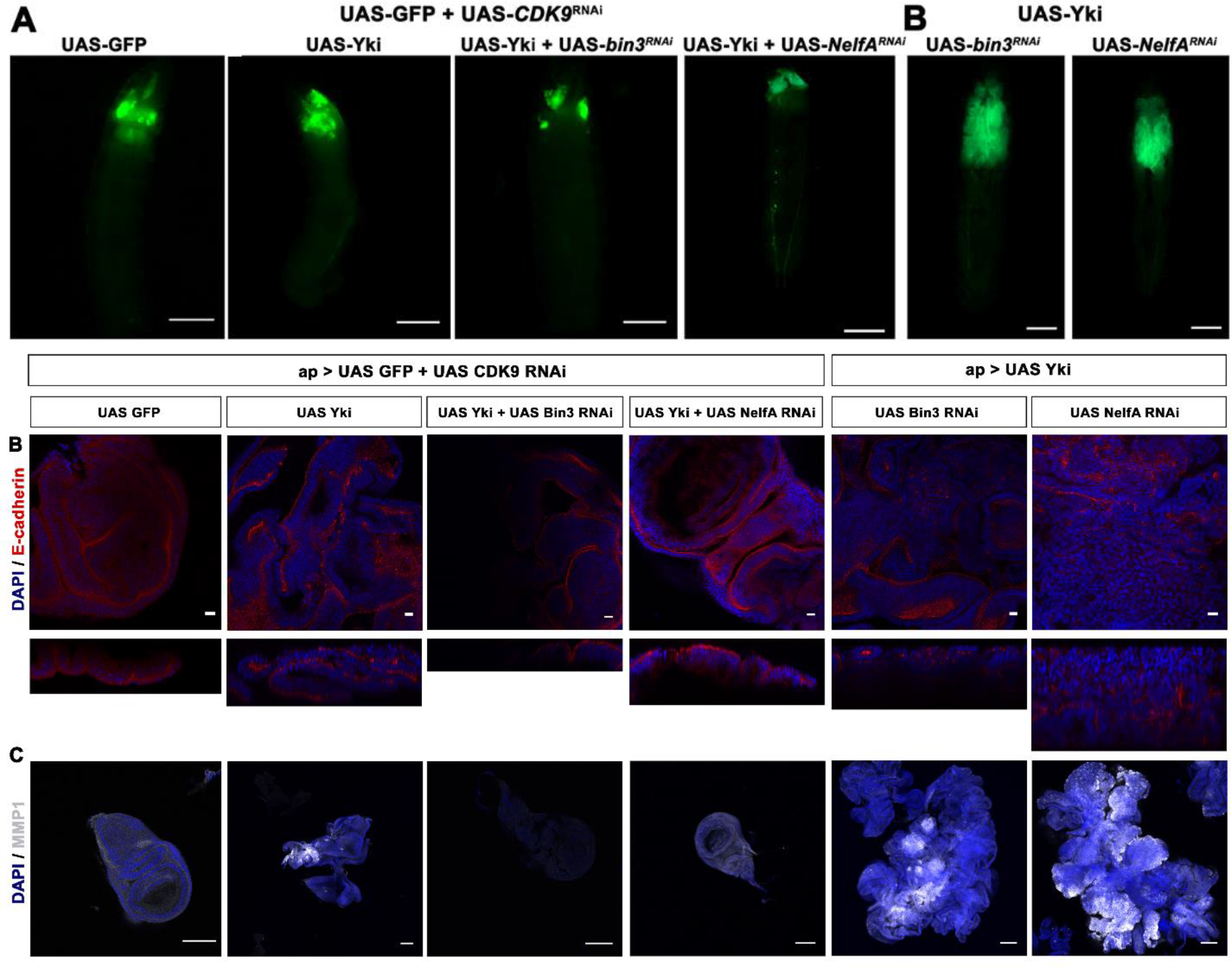
CDK9 is necessary for Yki-mediated tumorigenesis. A. Loss of CDK9 rescues tumor phenotype. The images show GFP-expressing wing discs of various genotypes as indicated. Size of the wing discs may be discerned by the amount of larval space occupied by GFP-expressing tissue. RNAi-mediated depletion of *cdk9* inhibited tumor formation caused by a combination of over-expression of Yki and depletion of a component of the PPP. The GFP-marked wing tissue is of the same size as in controls. All crosses were using GAL80^TS^; *ap*-GAL4; UAS-GFP (scale bar = 1 inch). B. Restoration of apico-basal polarity in wing disc tissue. The images show wing discs of various genotypes as indicated stained for E-Cad (white). RNAi-mediated depletion of *cdk9* restored normal apical localization of E-cad in wing discs that over-express Yki and also depleted for a component of the PPP. All discs are also stained with DAPI (blue) to visualize nuclei (scale bar = 10μm). C. Restoration of MMP1 levels. The images show wing discs of various genotypes as indicated stained MMP1 (white). RNAi-mediated depletion of *cdk9* restored normal levels of MMP1 in wing discs that over-express Yki and also depleted for a component of the PPP. All discs are also stained with DAPI (blue) to visualize nuclei (scale bar = 100μm).

Given that CDK9 is known to act directly on both NELF proteins and RNA Pol II, we wondered whether CDK9 activity would be required in the absence of the NELF complex. As shown above in the case of removing the 7SK snRNP complex, depletion of *cdk9* suppressed overgrowth caused by RNAi-mediated depletion of *NelfA* and over-expression of Yki (Fig. 4A). This was accompanied by restoration of apico-basal polarity and MMP1 expression to wild type levels (Fig 4B, C). This finding provides evidence that alleviation of pausing by removal of NELF complex is not sufficient without CDK9 activity. This presumably reflects an importance of activation of RNA Pol II by CDK9-mediated phosphorylation.

We then examined if depletion of the complexes associated with PPP and increased CDK9 levels are sufficient to cause over-growth phenotype or the growth is tightly coupled to the presence of a growth signal such as Yki. Depletion of components of 7SK snRNP or NELF complexes in the background of over-expressed CDK9 did not cause any growth phenotype or morphological alteration in wing disc epithelium (Fig. 5A). This suggests that deregulation of RNA Pol II pausing is not sufficient on its own to produce an over-growth or neoplastic phenotype; yet it does so in the context of Yki over-expression. In the context of elevated Yki activity, there appear to be two brakes, each of which must be removed by CDK9 activity to allow excess Yki to produce tumors in *Drosophila* wing disc tissue.

**Figure 5.**
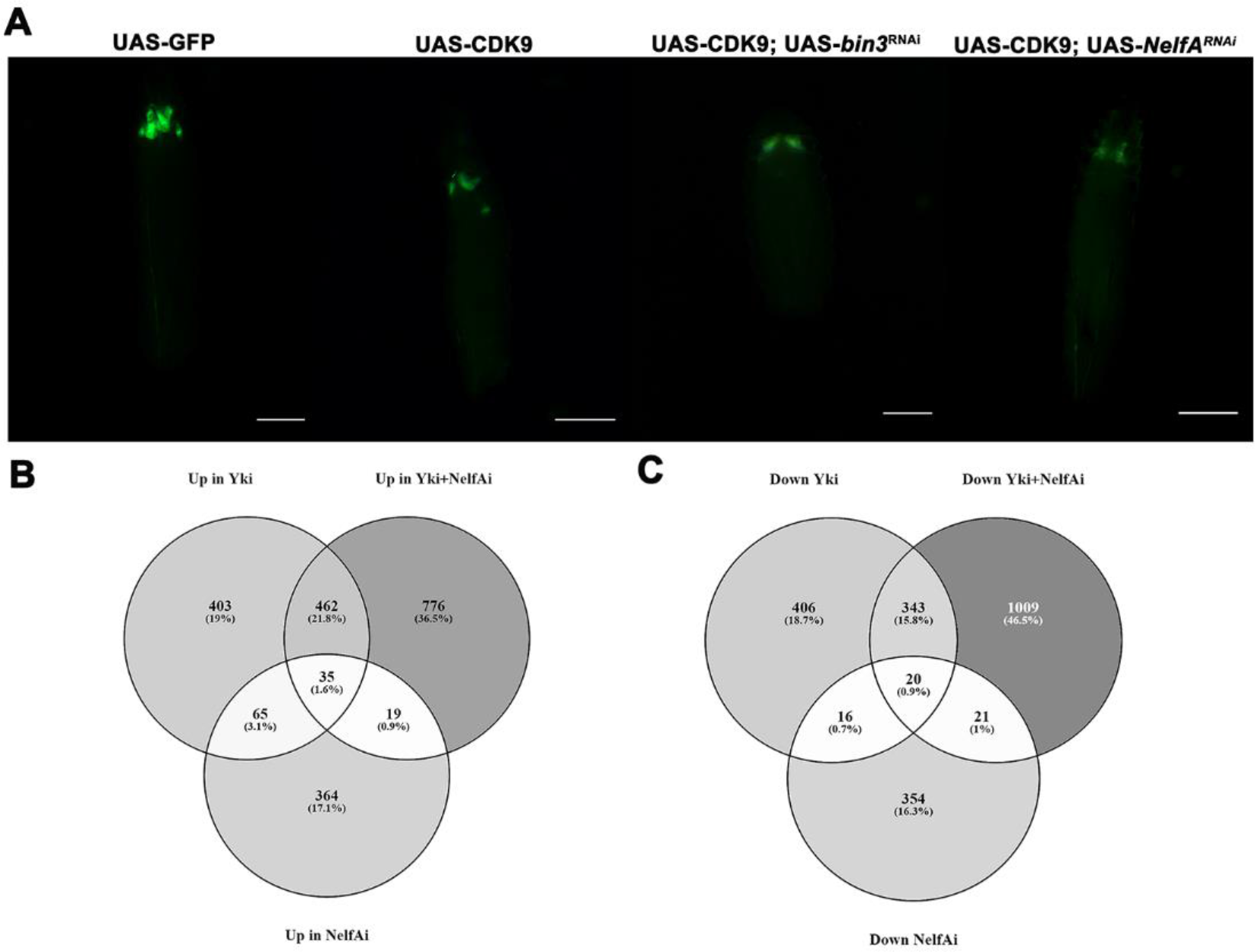
Yki is the driver of tumorigenesis. A. Larval images showing phenotype of UAS GFP in combination with (left to right) UAS-GFP, UAS-CDK9 followed by UAS-CDK9 and UAS-*bin3*^RNAi^, UAS-CDK9 and UAS-*NelfA*^RNAi^. None of them show over-growth phenotype as observed when Yki is over-expressed suggesting CDK9 may induce tumorous growth only in the context of over-expressed Yki. All crosses were using GAL80^TS^; *ap*-GAL4; UAS-GFP (scale bar = 1 inch). B. Venn diagram showing number of common and unique genes, who expression is upregulated in comparison with *ap* > *GFP* from different genotypes as indicated in the figure. C. Venn diagram depicting number of common and unique genes downregulated in comparison with *ap* > *GFP* from different genotypes.

### Tumorigenesis induced by alleviation of pausing is associated with deregulated proteostasis

As over-expression Yki was essential, although not sufficient, to cause neoplastic tumors, genetic experiments above provided an opportunity to distinguish between Yki activated genes that cause simple hyperplastic growth of the discs (when Yki is over-expressed in wild type background) vs to cause neoplastic growth (when Yki is over-expressed along with depletion of *bin3*, *Hexim* or NELFs).

We carried out RNA-seq to identify differentially expressed genes in tissues depleted for *NelfA* and overexpressing Yki as well as both individual treatments. We also carried out RNA-seq for GFP expressing wild type wing discs as control. Yki has previously been shown to regulate a number of transcripts involved in control of tissue growth and cell survival, including *bantam, DIAP1, cycE*. As expected, these transcripts were upregulated in the tumorous tissue. We find that 776 transcripts were uniquely upregulated (Fig. 5B) and 1009 transcripts were uniquely downregulated (Fig. 5C) in the tumorous wing discs, compared to all other genotypes including wildtype discs.

We also observed an enhancement of effect of Yki in a subset of transcripts that were common to non-tumorous tissue over-expressing Yki alone and tumorous tissue, wherein *NeflA* is depleted in the background of Yki over-expression. We reasoned that since PPP functions to attenuate expression of genes, the enhanced up- and down-regulation from common set of transcripts should be included in further analysis. Thus, we included 155 upregulated and 160 downregulated transcripts which were further up or down regulated in tumor tissue compared to Yki expressing tissue. We used this set of genes to do gene ontology (GO) analysis in order to explore gene sets that show enrichment and might indicate pathways or processes that are involved in tumorigenesis. For this purpose STRING tool was utilized [27]. STRING output is based on statistical enrichment score of interactions obtained from the input compared to a random set of genes from the genome of the organism in this case *D melanogaster*. STRING also collates data from manually curated databases of interactions such as KEGG and GO terms.

We observed enrichment for pathways involved in protein processing in endoplasmic reticulum, regulators of proteasome function as well as different components of proteasome among genes downregulated in tumorous tissues (Table 1). Interestingly, the upregulated set showed enrichment for ribosome and its biogenesis (Table 2). These observations indicate overall deregulation of protein homeostasis (proteostasis) in tumors caused by depletion of *NelfA* in combination with Yki overexpression, consistent with a recent data on human cancers [28,29].

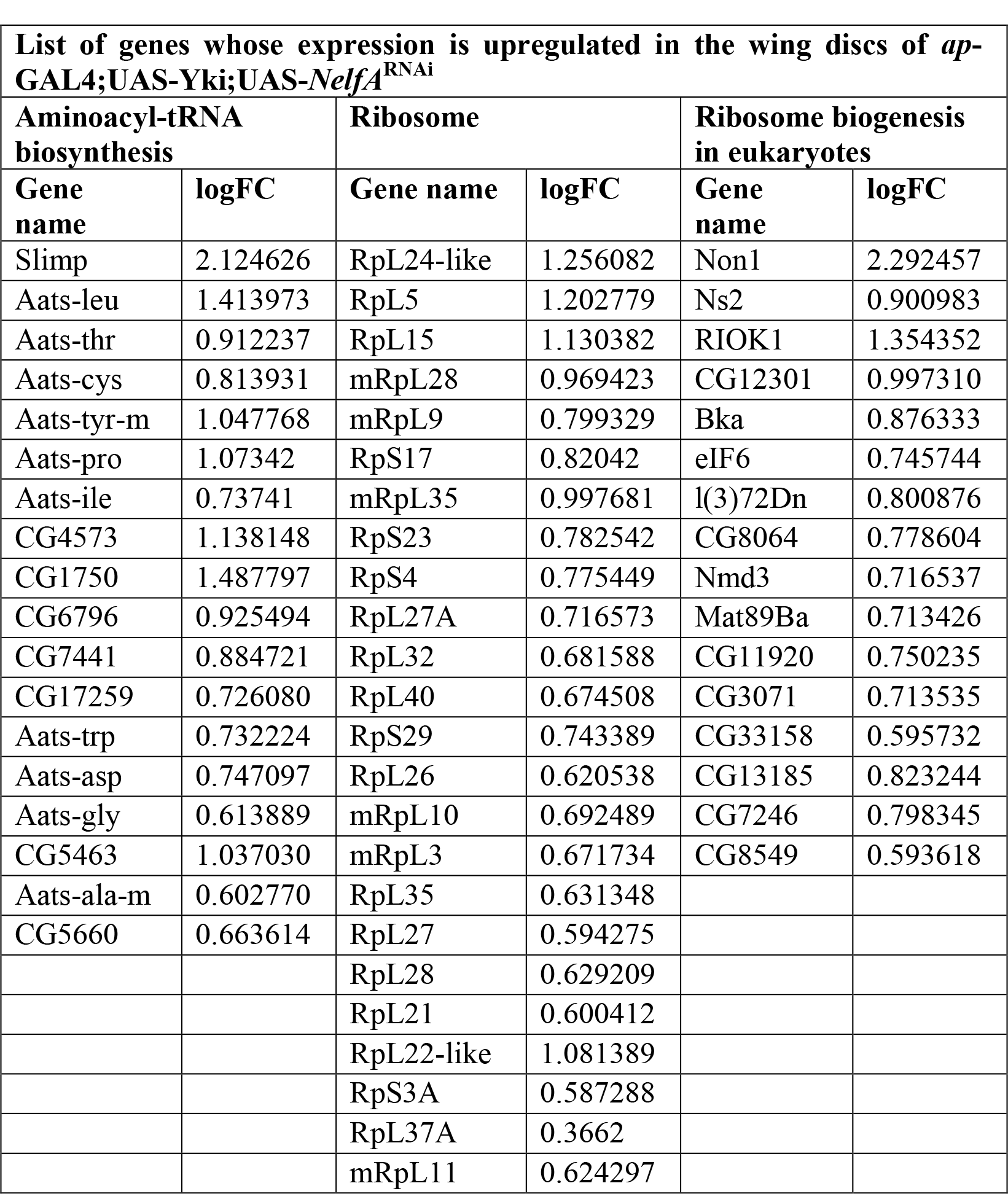

**Table 2**
| **List of genes whose expression is down regulated in the wing discs of *ap-GAL4;UAS-Yki;UAS-NelfA*<sup>RNAi</sup>** |  |  |  |
| --- | --- | --- | --- |
| **Proteasome** |  | **Protein processing in endoplasmic reticulum** |  |
| **Gene name** | **logFC** | **Gene name** | **logFC** |
| Rpn7 | -1.084698 |  |  |
| Rpn13 | -0.949757 | prtp | -1.46289 |
| Rpn2 | -0.900781 | Sec61gamma | -1.65593 |
| Prosalpha3 | -0.910536 | Sec61beta | -1.31964 |
| Rpn3 | -0.850895 | CG5885 | -1.28449 |
| Rpn1 | -0.879125 | Sec61alpha | -1.25184 |
| Pomp | -0.795409 | TRAM | -1.3381 |
| Prosalpha5 | -0.834712 | Pdi | -1.145 |
| Prosbeta4 | -0.76685 | SsRbeta | -1.23571 |
| Prosbeta7 | -0.717883 | Sec13 | -1.01315 |
| Prosbeta2 | -0.706638 | Sec63 | -0.97489 |
| Prosbeta5 | -0.687177 | CG14476 | -1.03296 |
| Prosalpha4 | -0.694159 | Sec24CD | -0.87522 |
| Prosbeta6 | -0.631819 | Ostgamma | -0.97374 |
| Rpn10 | -0.592037 | Ost48 | -0.88256 |
| Rpn12 | -0.597364 | CG4164 | -1.21065 |
|  |  | ergic53 | -0.86625 |
|  |  | Plap | -0.86843 |
|  |  | OstStt3 | -0.90274 |
|  |  | Gp93 | -0.89197 |
|  |  | l(1)G0320 | -0.88951 |
|  |  | CG33303 | -0.8065 |
|  |  | Hsc70-3 | -0.9493 |
|  |  | CG5510 | -0.81474 |
|  |  | p47 | -0.78553 |
|  |  | Crc | -0.86323 |
|  |  | CG6453 | -0.81869 |
|  |  | Sec23 | -0.73903 |
|  |  | ERp60 | -0.76882 |
|  |  | Der-1 | -0.80252 |
|  |  | Csp | -0.64369 |
|  |  | CaBP1 | -0.61193 |
|  |  | CG1597 | -0.67306 |

## Discussion

PPP has emerged as a critical regulatory step in gene expression [30]. It involves stalling of RNA Pol II in 20 to 60 nucleotides downstream of the transcription start site and controlled release of RNA Pol II when triggered by signals from the surroundings. Many studies in recent years have elucidated mechanisms by which RNA Pol II is stalled and the factors that bring about pausing as well as release of the paused RNA Pol II. Our *in vivo* model for tumorigenesis has allowed us to elucidate the functions of the NELF, 7SKsnRNP and P-TEFb complexes in the context of growth control *in vivo*. Previous studies have implicated NELF in regulating the response of embryonic stem cells to signaling cues such as FGF [31]. Furthermore, PPP has been shown to be important for coordination of expression genes involved in morphogenesis of *Drosophila* embryo [32]. Our findings provide direct evidence that PPP can limit tumor formation in the context of Hippo tumor suppressor pathway. Depletion of these factors alone or even in combination with over-expression of CDK9 was not sufficient to induce tumorous growth but did so when combined with over-expression of Yki. This cooperation appears to be specific to Yki induced tumors as there was no cooperation with other oncogenic drivers such as EGFR or activated Notch in wing disc tumor models. This suggests that pausing plays a previously unappreciated role regulating the output of Hippo pathway in growth control, thereby limiting its tumorigenic potential.

We were intrigued by the finding that CDK9 activity is required for Yki driven tumor formation, even when the upstream and downstream pausing complex factors have been removed. These observations suggest that CDK9 activity is required not only to remove the ‘brake’ exerted by NELF pausing complex, but also required to activate RNA Pol II through direct phosphorylation. Neither alone is sufficient. This suggests an overlapping ‘belt and suspenders’ regulation to ensure that expression of Yki targets is maintained at appropriate levels for normal growth control, while preventing over expression, which may lead to tumorigenesis. A mechanism of this sort allows for the possibility that other growth regulatory or metabolic homeostasis pathways might impact on the outcome of Yki activity via regulation of the CDK9. It will be of interest to explore how the interplay between these mechanisms is regulated during normal tissue growth. This also suggests a possible role for CDK9 inhibitors as a therapeutic approach to tumors related to activity of Hippo pathway.

Our genetic model is also useful to study the importance of PPP in attenuating transcriptional output at genome wide scale. Preliminary observations of data generated by RNA-seq suggest that most genes that are differentially expressed when Yki is overexpressed show further changes in the same direction (up or down regulation) in combination of Yki overexpression with depletion of Nelf-A. Furthermore, we also report deregulation of proteostasis uniquely in tumor tissue. This is consistent with recent reports that deregulation of translation and deregulation of protein processing are important factors in progression of cancers and might be target for therapy [28,29].

To conclude, our study has highlighted additional regulatory module on Yki driven tumorigenic activity which directly impinges on transcription. It will be interesting to see role of PPP machinery, which has been reported to be highly conserved from *Drosophila* to humans [33], in the context of highly conserved Hippo pathway effectors YAP/TAZ. Considering reported function of CDK9 in YAP driven transcription and therapeutic accessibility of CDK9 activity [20,34], it is critical to understand the function of 7SK snRNP and NELF complexes in this context.

## Material and methods

### *Drosophila* strains

Following *Drosophila* strains are used in this study. *ap-Gal4* [35], *UAS-Yki* [3]. Following RNAi stocks were obtained from Vienna *Drosophila* RNAi Center and Bloomington *Drosophila* stock Center: *UAS-NelfA*^*RNAi*^ (KK106245, TRiP #32897), *UAS-NelfB*^*RNAi*^ (KK108441, TRiP #42547), *UAS-NelfE*^*RNAi*^ (TRiP # 32835), *UAS-NelfD*^*RNAi*^ (KK100009, TRiP # 38934, #42931), *UAS*-*bin3*^*RNAi*^ (KK101090, TRiP #41527), *UAS-Hexim*^*RNAi*^ (KK100500). *UAS-CDK9* was obtained from FlyORF (#F001571).

### Spatio-temporal regulation of transgene expression in wing imaginal disc

*apterous* enhancer was used to drive expression of *Gal4* conditionally in dorsal compartment of wing imaginal discs. *Gal4* activity was regulated using Gal80^TS^ which allows expression of transgenes at permissive temperature of 29° C as against restrictive 18° C temperature. In all experiments, *tubulin-*Gal80^TS^ was used. *Drosophila* crosses were allowed to lay eggs for three days at 18° C and were then flipped or discarded. Larvae were then allowed to grow for additional 5 days before switching to temperature of 29° C. At 29° C they were maintained for 4-14 days. All crosses were using *tubulin*-GAL80^TS^; *ap*-GAL4; UAS-GFP. Thus, all experimental crosses had one copy of GFP, while control crosses had two copies of GFP.

### Immunohistochemistry

Following primary antibodies were used: rat anti-Ecadherin, mouse anti-MMP1 (Developmental Studies Hybridoma Bank). Rhodamine-phalloidin (ThermoFisher Scientific,Cat no R415) was used to stain actin in tissue.

3^rd^ instar larvae were dissected in PBS. Samples were fixed in 4% PFA for 20 minutes, followed by three 10 minutes washes in PBT (PBS-Tween20) at room temperature. 5% BSA in PBS was used for blocking followed by overnight incubation in primary antibody at 4° C. Next day the samples were washed with PBT, three times for 10 minutes each followed by incubation with secondary antibody for two hours at room temperature. Samples were then washed with PBT and stained for DNA using DAPI (Sigma Aldrich) for 5 min. Wing disc tissue was then mounted on slides in Antifade Gold mountant (ThermoFisher Scientific). Imaging was done on Leica SP8 confocal laser-scanning microscope. Image processing was done using ImageJ and Adobe Photoshop 6.

### Whole Larval imaging

Larval images were taken in bright field and in GFP channel with Leica stereomicroscope. Image processing was done using Adobe Photoshop 6 and ImegeJ.

### RNA-seq

Induction procedure for transgenes was followed as mentioned earlier. Wing imaginal disc tissue was dissected on 4-5^th^ day after induction for *ap*>*GFP*, *ap*>*UAS-Yki*, *ap*>*Nelf-A RNAi* (KK106245), *ap*>*UAS-Yki*, *UAS Nelf-A RNAi*. Larvae were washed in RNase, DNase free ultrapure water (GiBCO) and then dissections were done in RNase, DNase free PBS (GiBCo). Number of wing imaginal discs collected was 150, 70, 150, 25 respectively for *ap*>*GFP*, *ap*>*UAS-Yki*, *ap*>*Nelf-A RNAi* (KK106245), *ap*>*UAS-Yki*, *UAS Nelf-A RNAi*. Collection was done in TRiZOL reagent (ThermoFisher Scientific). Each genotype was collected in three biological replicates. RNA sequencing was done on Illumina platform.

### RNA-seq data analysis

RNA-seq analysis was performed using the HISAT 2.0 package protocol as explained in Pertea *et al.*, 2016. To identify significantly differentially expressing genes in different combinations of comparisons, DEseq package and EdgeR were used [36].

The list of genes obtained was then used as an input for web based tool venny (Oliveros, J.C. (2007) VENNY. An interactive tool for comparing lists with Venn Diagrams. http://bioinfogp.cnb.csic.es/tools/venny/index.html) to obtain list of genes that are unique to each genotype, overlapping between all three or combination of any two genotypes.

### Gene ontology (GO) analysis

For GO and pathway enrichment analysis, we utilized STRING10 [27]. We used gene lists that are significantly differentially expressed in single genotype or a combination of genotypes as mentioned in the results section, as input to the STRING. The output files were downloaded as interaction network and list of genes from input that are enriched in different GO categories or as KEGG pathways.

## Acknowledgements

This work was primarily supported by an Indo-Danish research grant from Department of Biotechnology, Govt. of India to LSS and from Innovation fund Denmark, Novo Nordisk Foundation NNF12OC0000552 and Neye Foundation to SMC. JC Bose Fellowship and grant from Department of Science & Technology, Govt of India to LSS. UGC Research Fellowship to SN. We thank other members of both the laboratories for critical input.

## Author Contributions

SN, PG and RW carried out all fly experiments. SN did RNA-seq and its analysis, all image analyses and wrote the MS. LSS and SM conceived the project and wrote the MS.

## Expanded View Figures

**Fig EV1.**
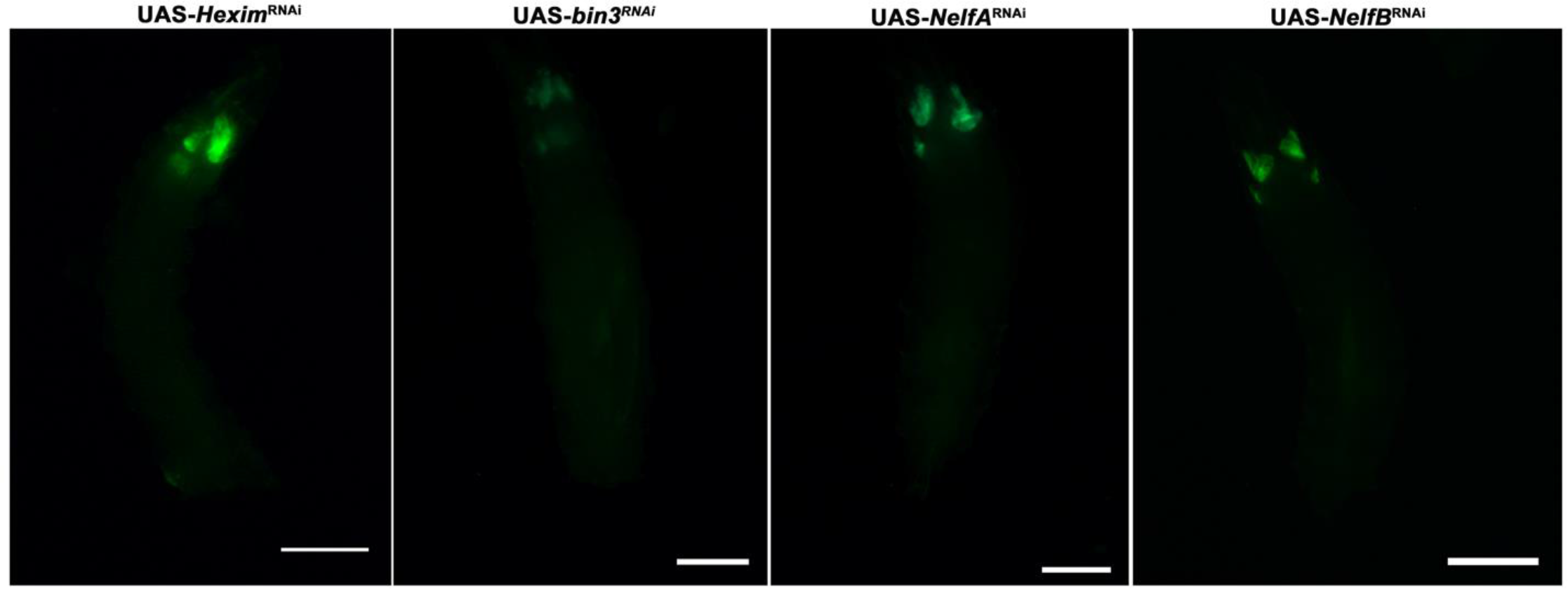
The images show GFP-expressing wing discs of various genotypes as indicated. Down regulation of the any of the components of 7skRNP or NEFL complexes on their own do not cause any over-growth phenotype. All crosses were using GAL80^TS^; *ap*-GAL4; UAS-GFP.

**Fig EV2.**
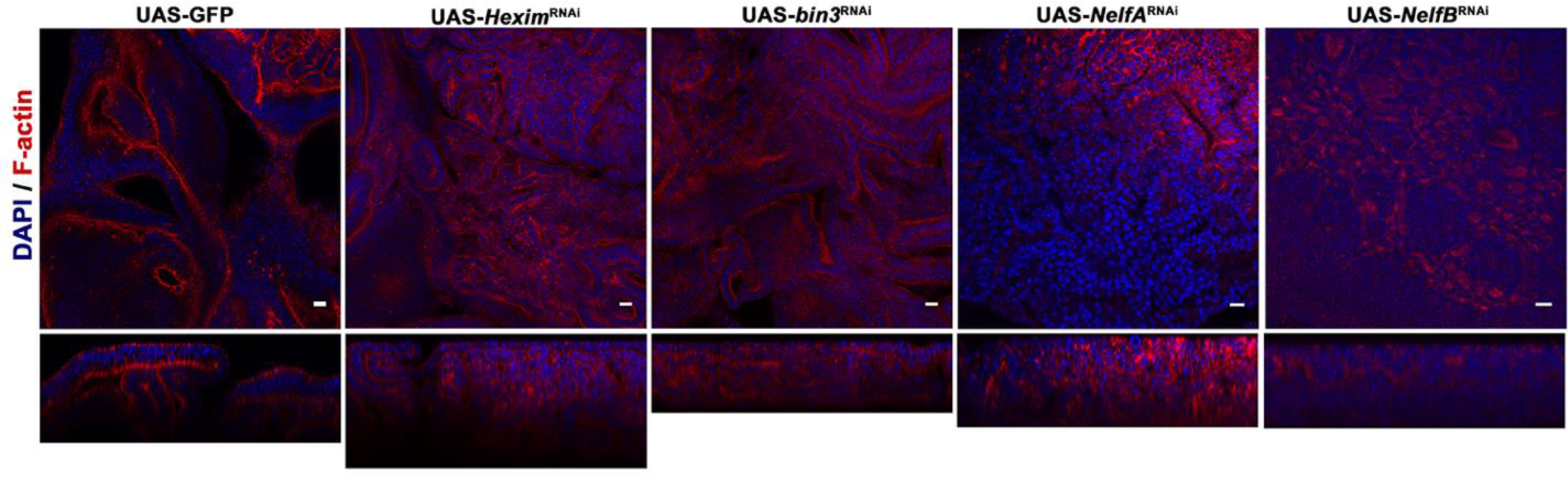
Disruption of characteristic epithelial apico-basal polarity in tumor discs. Images of wing discs over-expressing Yki alone (crossed to UAS-GFP as control) or in combination RNAi-mediated knowckdown of *Hexim, bin3, NelfA* or *NelfB*. All crosses were using GAL80^TS^; *ap*-GAL4; UAS-GFP. Discs are stained with Phalloidin (red), which stain F-Actin. expression and localization. Bottom panel of each image shows orthogonal optical section of respective genotype. Note delocalization of F-actin in tumorous tissues caused by the depletion of a component of PPP and Yki over-expression. All discs are also stained with DAPI (blue) to visualize nuclei.

## References

1. Pan D (2010) The hippo signaling pathway in development and cancer. Dev Cell 19: 491–505.

2. Zhao B, Tumaneng K, Guan KL (2011) The Hippo pathway in organ size control, tissue regeneration and stem cell self-renewal. Nat Cell Biol 13: 877–883.

3. Huang J, Wu S, Barrera J, Matthews K, Pan D (2005) The Hippo signaling pathway coordinately regulates cell proliferation and apoptosis by inactivating Yorkie, the Drosophila homolog of YAP. Cell 122: 421–434.

4. Dong J, Feldmann G, Huang J, Wu S, Zhang N, Comerford SA, Gayyed MF, Anders RA, Maitra A, Pan D (2007) Elucidation of a Universal Size-Control Mechanism in Drosophila and Mammals. Cell 130: 1120–1133.

5. Zanconato F, Forcato M, Battilana G, Azzolin L, Quaranta E, Bodega B, Rosato A, Bicciato S, Cordenonsi M, Piccolo S (2015) Genome-wide association between YAP/TAZ/TEAD and AP-1 at enhancers drives oncogenic growth. Nat Cell Biol 17:.

6. Zender L, Spector MS, Xue W, Flemming P, Cordon-Cardo C, Silke J, Fan ST, Luk JM, Wigler M, Hannon GJ, et al. (2006) Identification and Validation of Oncogenes in Liver Cancer Using an Integrative Oncogenomic Approach. Cell 125: 1253–1267.

7. Overholtzer M, Zhang J, Smolen GA, Muir B, Li W, Sgroi DC, Deng C-X, Brugge JS, Haber DA (2006) Transforming properties of YAP, a candidate oncogene on the chromosome 11q22 amplicon. Proc Natl Acad Sci U S A 103: 12405–12410.

8. Johnson R, Halder G (2014) The two faces of Hippo: Targeting the Hippo pathway for regenerative medicine and cancer treatment. Nat Rev Drug Discov 13: 63–79.

9. Groth C, Vaid P, Khatpe A, Prashali N, Ahiya A, Andrejeva D (2019) Genome-wide RNAi screen for context-dependent tumor suppressors identified using in vivo models for neoplasia in *Drosophila*. BioRxiv preprint: https://doi.org/10.1101/643221.

10. Guenther MG, Levine SS, Boyer LA, Jaenisch R, Young RA (2007) A Chromatin Landmark and Transcription Initiation at Most Promoters in Human Cells. Cell 130: 77–88.

11. Muse GW, Gilchrist DA, Nechaev S, Shah R, Parker JS, Grissom SF, Zeitlinger J, Adelman K (2007) RNA polymerase is poised for activation across the genome. Nat Genet 39: 1507–1511.

12. Zeitlinger J, Stark A, Kellis M, Hong J-WW, Nechaev S, Adelman K, Levine M, Young RA (2007) RNA polymerase stalling at developmental control genes in the Drosophila melanogaster embryo. Nat Genet 39: 1512–1516.

13. Kwak H, Lis JT (2013) Control of Transcriptional Elongation. Annu Rev Genet 47: 483–508.

14. Yamaguchi YY, Takagi TT, Wada TT, Yano KK, Furuya AA, Sugimoto SS, Hasegawa JJ, Handa HH (1999) NELF, a multisubunit complex containing RD, cooperates with DSIF to repress RNA polymerase II elongation. Cell 97: 11.

15. Wada T, Takagi T, Yamaguchi Y, Ferdous A, Imai T, Hirose S, Sugimoto S, Yano K, Hartzog GA, Winston F, et al. (1998) DSIF, a novel transcription elongation factor that regulates RNA polymerase II processivity, is composed of human Spt4 and Spt5 homologs. Genes Dev 12: 343–356.

16. Marshall NF, Price DH (1995) Purification of P-TEFb, a transcription factor required for the transition into productive elongation. J Biol Chem 270: 12335–12338.

17. Jonkers I, Lis JT (2015) Getting up to speed with transcription elongation by RNA polymerase II. Nat Rev Mol Cell Biol 16: 167–177.

18. Nguyen D, Krueger BJ, Sedore SC, Brogie JE, Rogers JT, Rajendra TK, Saunders A, Matera AG, Lis JT, Uguen P, et al. (2012) The Drosophila 7SK snRNP and the essential role of dHEXIM in development. Nucleic Acids Res 40: 5283–5297.

19. Kanazawa S, Soucek L, Evan G, Okamoto T, Peterlin BM (2003) c-Myc recruits P-TEFb for transcription, cellular proliferation and apoptosis. Oncogene 22: 5707–5711.

20. Galli GG, Carrara M, Yuan WC, Valdes-Quezada C, Gurung B, Pepe-Mooney B, Zhang T, Geeven G, Gray NS, de Laat W, et al. (2015) YAP Drives Growth by Controlling Transcriptional Pause Release from Dynamic Enhancers. Mol Cell 60: 328–337.

21. Gateff E, Löffler T, Wismar J (1993) A temperature-sensitive brain tumor suppressor mutation of Drosophila melanogaster: Developmental studies and molecular localization of the gene. Mech Dev 41: 15–31.

22. Tanos B, Rodriguez-Boulan E (2008) The epithelial polarity program: Machineries involved and their hijacking by cancer. Oncogene 27: 6939–6957.

23. Beaucher M, Hersperger E, Page-McCaw A, Shearn A (2007) Metastatic ability of Drosophila tumors depends on MMP activity. Dev Biol 303: 625–634.

24. Uhlirova M, Bohmann D (2006) JNK- and Fos-regulated Mmp1 expression cooperates with Ras to induce invasive tumors in Drosophila. EMBO J 25: 5294–5304.

25. Miles WO, Dyson NJ, Walker J a. (2011) Modeling tumor invasion and metastasis in Drosophila. Dis Model Mech 4: 753–761.

26. Jennings BH (2013) Pausing for thought: Disrupting the early transcription elongation checkpoint leads to developmental defects and tumourigenesis. BioEssays 35: 553–560.

27. Szklarczyk D, Morris JH, Cook H, Kuhn M, Wyder S, Simonovic M, Santos A, Doncheva NT, Roth A, Bork P, et al. (2017) The STRING database in 2017: Quality-controlled protein-protein association networks, made broadly accessible. Nucleic Acids Res 45: D362–D368.

28. Ruggero D (2013) Translational control in cancer etiology. Cold Spring Harb Perspect Biol 5:.

29. Pelletier J, Thomas G, Volarevic S, Volarević S (2017) Nrc.2017.104. Nature 18: 51.

30. Core L, Adelman K (2019) polymerase II : a nexus of gene regulation. 1–24.

31. Williams LH, Fromm G, Gokey NG, Henriques T, Muse GW, Burkholder A, Fargo DC, Hu G, Adelman K (2015) Pausing of RNA Polymerase II Regulates Mammalian Developmental Potential through Control of Signaling Networks. Mol Cell 58: 311–322.

32. Lagha M, Bothma JP, Esposito E, Ng S, Stefanik L, Tsui C, Johnston J, Chen K, Gilmour DS, Zeitlinger J, et al. (2013) XPaused Pol II coordinates tissue morphogenesis in the drosophila embryo. Cell 153: 976.

33. Peterlin BM, Price DH (2006) Controlling the Elongation Phase of Transcription with P-TEFb. Mol Cell 23: 297–305.

34. Blake DR, Vaseva A V., Hodge RG, Kline MP, Gilbert TSK, Tyagi V, Huang D, Whiten GC, Larson JE, Wang X, et al. (2019) Application of a MYC degradation screen identifies sensitivity to CDK9 inhibitors in KRAS-mutant pancreatic cancer. Sci Signal 12: eaav7259.

35. Cohen B, McGuffin ME, Pfeifle C, Segal D, Cohen SM (1992) apterous, a gene required for imaginal disc development in Drosophila encodes a member of the LIM family of developmental regulatory proteins. Genes Dev 6: 715–729.

36. Anders S, Huber W (2010) Differential expression analysis for sequence count data. Genome Biol 11: R106.

